# Protection conferred by intraperitoneal Group A *Streptococcus* immunization relies on macrophages and IFN-γ but not on concurrent adaptive immune responses

**DOI:** 10.1101/2023.10.31.564887

**Authors:** Shiva Emami, Elsa Westerlund, Thiago Rojas Converso, Bengt Johansson-Lindbom, Jenny J Persson

## Abstract

Group A *Streptococcus* (GAS; *Streptococcus pyogenes*) is an important bacterial pathogen estimated to cause over 700 million superficial infections and around 500.000 deaths due to invasive disease or severe post-infection sequelae in the world yearly. In spite of this major impact on society, there is currently no vaccine available against this bacterium. GAS strains can be separated into >200 distinct *emm* (M)-types, and protective immunity against GAS is believed to in part be dependent on type-specific antibodies. Here, we analyze the nature of protective immunity generated against GAS in a model of intraperitoneal immunization in mice. We demonstrate that multiple immunizations are required for the ability to survive a subsequent lethal challenge, and although significant levels of GAS-specific antibodies are produced, these are redundant for protection. Instead, our data show that the immunization-dependent protection in this model is induced in the absence of B and T cells, is accompanied by an altered cytokine profile upon subsequent infection and requires macrophages and the macrophage-activating cytokine IFN-γ. To our knowledge these findings are the first to suggest that GAS has the ability to induce forms of trained innate immunity. Taken together, the current study reveals a novel mechanism of the innate immune system in response to GAS infections that potentially could be leveraged for future development of effective vaccines.

**Author summary:** The bacterium Group A Streptococcus (GAS) causes many hundred million infections and around 500.000 deaths in the world every year. GAS can give rise to a wide spectrum of diseases ranging from mild strep throat to life-threatening necrotizing fasciitis (often referred to as “flesh-eating disease”). There is currently no vaccine available for this pathogen, much due to our incomplete knowledge of how the immune system reacts to different GAS infections, what immune responses are in fact required for long-term protection and how these are generated. Here we show that protective immunity arising after immunization through the intraperitoneal (ip) cavity requires multiple injections using heat killed GAS. Surprisingly, although typical adaptive immune responses are activated and generate production of GAS-specific antibodies these are redundant for protection, which instead hinges on macrophages and the cytokine IFN-γ. Our findings suggest that ip GAS immunizations trigger what is known as ‘trained immunity’, where innate immune cells become imprinted to respond with increased efficiency towards subsequent infection. Overall, these observations highlight a previously unknown ability of GAS to induce non-canonical forms of protective immunity, discoveries that may significantly contribute to our thinking about how the immune system reacts to such infections and broaden the scope for future vaccine strategies.

## Introduction

Group A *Streptococcus* (GAS, *Streptococcus pyogenes*) is one of our major circulating bacterial pathogens, causing >700 million uncomplicated infections and ∼500.000 deaths annually [1]. The vast majority of GAS infections are superficial (*e.g.* tonsillitis or impetigo) and occur in children, while uncommon devastating infections like necrotizing fasciitis or toxic shock syndrome occur predominantly in the adult or elderly population [1]. In rare cases, recurring and untreated mild infections may give rise to the autoimmune syndrome acute rheumatic fever (ARF), which may progress to life-threatening rheumatic heart disease [2, 3]. ARF is believed to at least in part be caused by immune responses generated against the GAS M protein that cross-react with *e.g.* cardiac myosin in the host [2]. The M protein is a major virulence factor and contains a hypervariable region (HVR) that allows for the differentiation of GAS strains into >220 *emm* types and is also target for protective, type-specific antibodies [4]. The extensive strain variability, immuno-subdominance of the HVR [5, 6] and the risk of vaccine components triggering ARF [7] are all major challenges in GAS vaccine development and there is currently no vaccine available against GAS.

Immunological memory is formed from highly specific B and/or T cells. During the resolution phase after an infection, expanded populations of specific B and T cells contract and leave behind memory cells with heightened activation potential. Recent advances however have shown that the spectrum of lasting infection-or vaccine-induced immune protection is broader than previously appreciated, and may include several forms of imprinted memory-like functions driven by metabolic and epigenetic changes in innate immune cell populations (trained immunity) [8]. In contrast to adaptive B and T cell memory, imprinted innate responses are non-specific and thus confer protection against both homologous and heterologous infections. Although our knowledge is clearly still limited, trained immunity has been extensively studied in myeloid populations like monocytes/macrophages and their precursors in the bone marrow, and has also been described in lymphoid NK cells, ILCs and γδ T cells [8].

Understanding the mechanisms underlying natural or experimentally induced immune protection against GAS has been a topic of interest for decades, and most attention has been devoted to the type-specific immunity GAS infections may confer [9–11]. In addition, it also seems that a more general protection may be present in adults as compared to children [12, 13] which to some degree might be explained by partial cross-protection observed within clusters of similar GAS strains building up over time [14, 15]. Whether or not trained immunity may contribute to the broader protection against GAS infection observed with age has not been investigated, and neither has the potential ability of GAS itself to induce innate memory forms.

In this study, we investigate protective immunity conferred by intraperitoneal (ip) immunization of mice with intact heat killed (HK) GAS. We find that repeated injections are required to establish protection against subsequent infection, and that protection induced through this immunization regimen seems to be independent of adaptive immunity. Instead, macrophages and the cytokine IFN-γ are essential for survival of infected immunized animals, indicating that GAS immunizations may generate a form of imprinted immunity in innate immune cell populations.

## Results

### Repeated intraperitoneal immunization with heat-killed GAS-M1 confers protection

The route of infection or vaccination strongly affects the quality and quantity of the initiated subsequent immune response, both in humans and in experimental murine models [16, 17]. In mice, intraperitoneal (ip) injections are commonly used both to model systemic infection, as they support rapid dissemination of the pathogen, and for immunization/vaccination regimens as they accommodate self-drain of antigen to the lymphatic system [18]. To identify an immunization strategy that would provide protection against a systemic (ip) GAS challenge, we first conducted a set of one-dose immunizations as follows: 1. 10^6^ CFU live GAS-M1 (highest possible non-lethal dose (data not shown)), 2. 10^8^ CFU formalin-fixed (FF) GAS-M1, and 3. 10^8^ CFU HK GAS-M1, were injected ip into C57Bl/6 (B6) mice. Control mice received PBS only. Three weeks after immunization, mice were challenged ip with a lethal dose of 10^8^ CFU GAS-M1 and monitored for moribundity. As neither of these immunizations provided protection above control levels (Fig 1A), we then designed an extended protocol employing the same immunization formulations (10^6^ live, or 10^8^ FF or HK GAS-M1) but using three injections with three-week intervals. In addition, all animals were bled 19 days after each immunization (Fig 1B). Three injections with live bacteria did still not provide increased protection as compared to control mice receiving PBS only during the immunization scheme. In contrast, three immunizations using FF or HK GAS-M1 both provided protection, and although not statistically significant, mice immunized with HK bacteria consistently displayed an increased survival as compared to mice immunized with FF bacteria (Fig 1C).

As antibodies commonly are important for protection against extracellular bacterial pathogens, we followed the serum GAS-M1-specific IgG levels through the immunization regimens. While live and FF bacteria did not stimulate any significant increases in GAS-M1-specific IgG even after three immunizations, HK bacteria induced significantly elevated levels after two immunizations (∼1.5 log) and a similar further increase after the third dose (Fig 1D). Protection induced by immunization with FF bacteria did therefore not seem to correlate with a robust increase in antibody production. In agreement with the lack of protection observed after the one-dose immunization regimens, none of the formulations promoted IgG generation after a single dose.

We were intrigued by the apparent inability of live bacteria to induce protection and measurable antibody responses but were unable to make proper comparisons to the FF or HK bacterial injections because of the differences in distributed dose (FF=HK: 10^8^, live: 10^6^). Therefore, we immunized mice with the same lower dose of HK and live GAS-M1 and observed that at this lower dose, HK GAS-M1 similarly failed to induce both protection and specific IgG after three immunizations (S1 Fig A and B). These findings show that the provided protection is dose dependent, and thus prevent us from drawing conclusions regarding the intrinsic ability or inability of live bacteria to confer protective immunity using this regimen. Based on these findings, we opted to use a dose of 10^8^ CFU of HK GAS-M1 for all subsequent immunization experiments.

**Fig 1.**
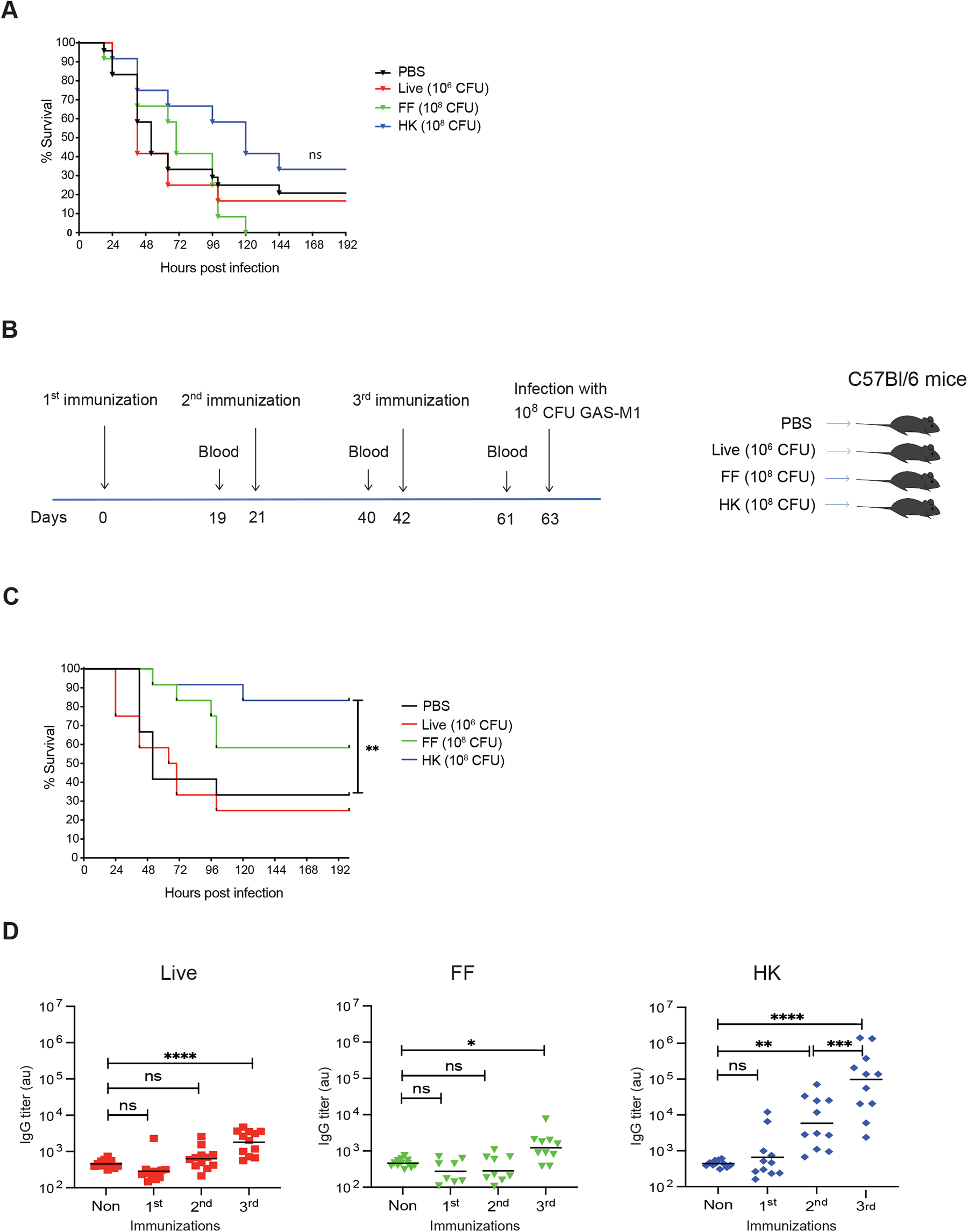
Protective immunity after repeated intraperitoneal immunization with HK GAS-M1. B6 mice (n= 12 per group) were immunized ip with one or three doses of HK GAS-M1 (10^8^ CFU), FF GAS-M1 (10^8^ CFU), or live GAS-M1 (10^6^ CFU) with three weeks interval between each dose. Control B6 mice received PBS only. Three weeks after completed immunization schedule, mice were challenged ip with a lethal dose of GAS-M1 (10^8^ CFU) and monitored for 8 days. (A) Kaplan-Meier survival plot with log-rank tests following immunization with one dose only. (B) Experimental layout for experiments based on three consecutive immunizations. (C) Kaplan-Meier survival plot with log-rank tests for mice immunized with three consecutive doses. (D) Serum level of IgG against intact GAS-M1 19 days after each dose of live, FF and HK GAS-M1. Statistical significance was analyzed by one-way ANOVA with Tukey’s multiple comparisons test. *p<0.05, **p<0.01, ***p<0.001 and ****p<0.0001. ns = not significant.

### Protective immunity after intraperitoneal immunization does not rely on germinal center-derived antibodies

High affinity and isotype switched antibody responses develop mostly within germinal centers (GC), histological structures appearing in B cell follicles of secondary lymphoid organs after infections or vaccination. GC B cell responses are dependent on T follicular helper (Tfh) cells, a subset of CD4^+^ T cells functionally specialized in providing B cell help [19]. Although isotype switching can occur also during T cell-independent B cell activation, these responses are mostly associated with production of low-affinity IgM [20]. To investigate the role for GC-derived antibodies in the protection acquired through ip immunization with HK GAS-M1, we employed the CD4-Cre.Bcl6^fl/fl^ mouse model. In this model, Cre^+^ mice lack the transcription factor Bcl6 in T cells, preventing Tfh cell differentiation and GC formation [21–23]. Surprisingly, we observed no significant difference in survival after infection (Fig 2A), or in IgG levels (Fig 2B) after 3 immunizations between animals lacking *Bcl6* in T cells and fully *Bcl6* competent animals. Further, GC-deficient animals did not exhibit a significant compensatory increase in specific antibodies of the complement-fixing IgM isotype that could explain the observed protection (Fig 2C). Taken together, our data indicate that GC formation and GC-derived antibodies are not essential for protection in this model, and that the generated antibodies are in fact not predominantly derived from GCs.

**Fig 2.**
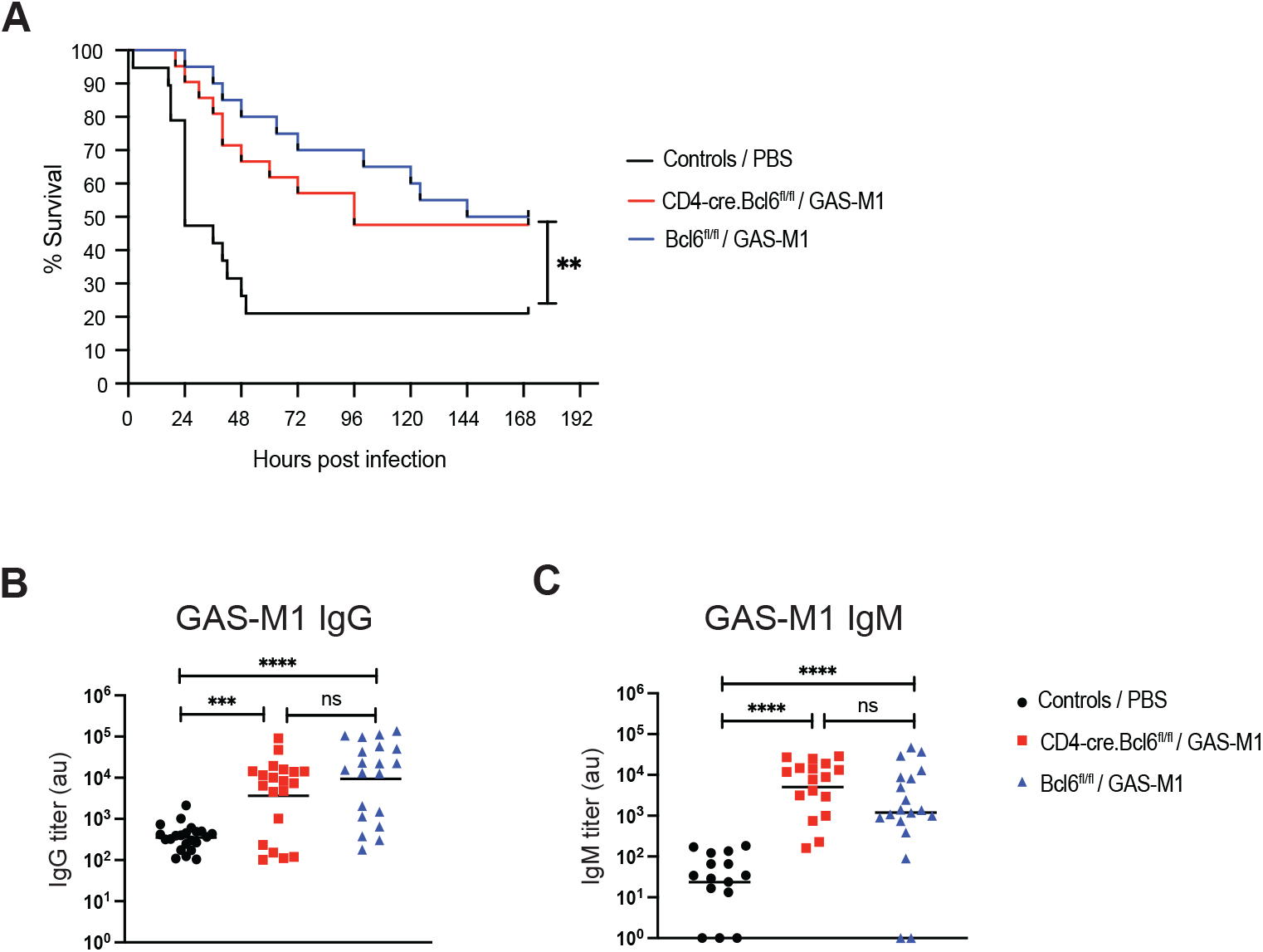
Protection conferred through the intraperitoneal immunization regimen does not depend on GC-derived antibodies. CD4-Cre.Bcl6^fl/fl^ (n=20) and Bcl6^fl/fl^ (n=21) mice received three ip immunization injections with HK GAS-M1. A control group (n=19), including mice of each genotype, received three doses of PBS only. Three weeks after the last dose, mice were challenged ip with 10^8^ CFU of live GAS-M1 and the animals were monitored for seven days post-infection. Pooled results from three separate experiments are shown. (A) Kaplan-Meier survival plot with log-rank tests. (B-C) Serum level of IgG (B) and IgM (C) against intact GAS-M1 19 days after the third immunization and two days prior to administration of the lethal dose. Statistical significance was analyzed by one-way ANOVA with Tukey’s multiple comparisons test. **p<0.01, ***p<0.001 and ****p<0.0001. ns = not significant.

### Protective immunity does not require T cell help during lethal GAS-M1 infection in immunized animals

In addition to providing help to B cells, memory Th cell subsets can facilitate clearance of bacteria through cytokine-mediated effects on myeloid cells, including macrophages and neutrophils [24]. To assess if the ip immunization regimen with HK GAS-M1 induces memory Th cells with such protective function, a depleting anti-CD4 monoclonal antibody (mAb) was used. One single dose of anti-CD4 mAb efficiently depleted CD4+ T cells, as assessed in spleen one and four days later, but did not reduce the percentage of CD8^+^ T cells (Fig 3A and B). We administered the anti-CD4 mAb to immunized and control mice 2 days prior to lethal infection and then every third day for the whole duration of the experiment. As shown in figure 3C, immunized CD4^+^ T cell-depleted animals were protected against the lethal dose to the same extent as non-depleted animals (Fig 3C). This was not due to inefficient depletion of CD4^+^ T cells, as demonstrated by the absence of CD4^+^ T cells in immunized animals surviving until the endpoint of the experiment (Fig 3D). While this experimental set-up does not exclude the requirement for T cell help during immunizations, we conclude that the anamnestic response that protect following administration of the lethal dose is independent of T cell help.

**Fig 3.**
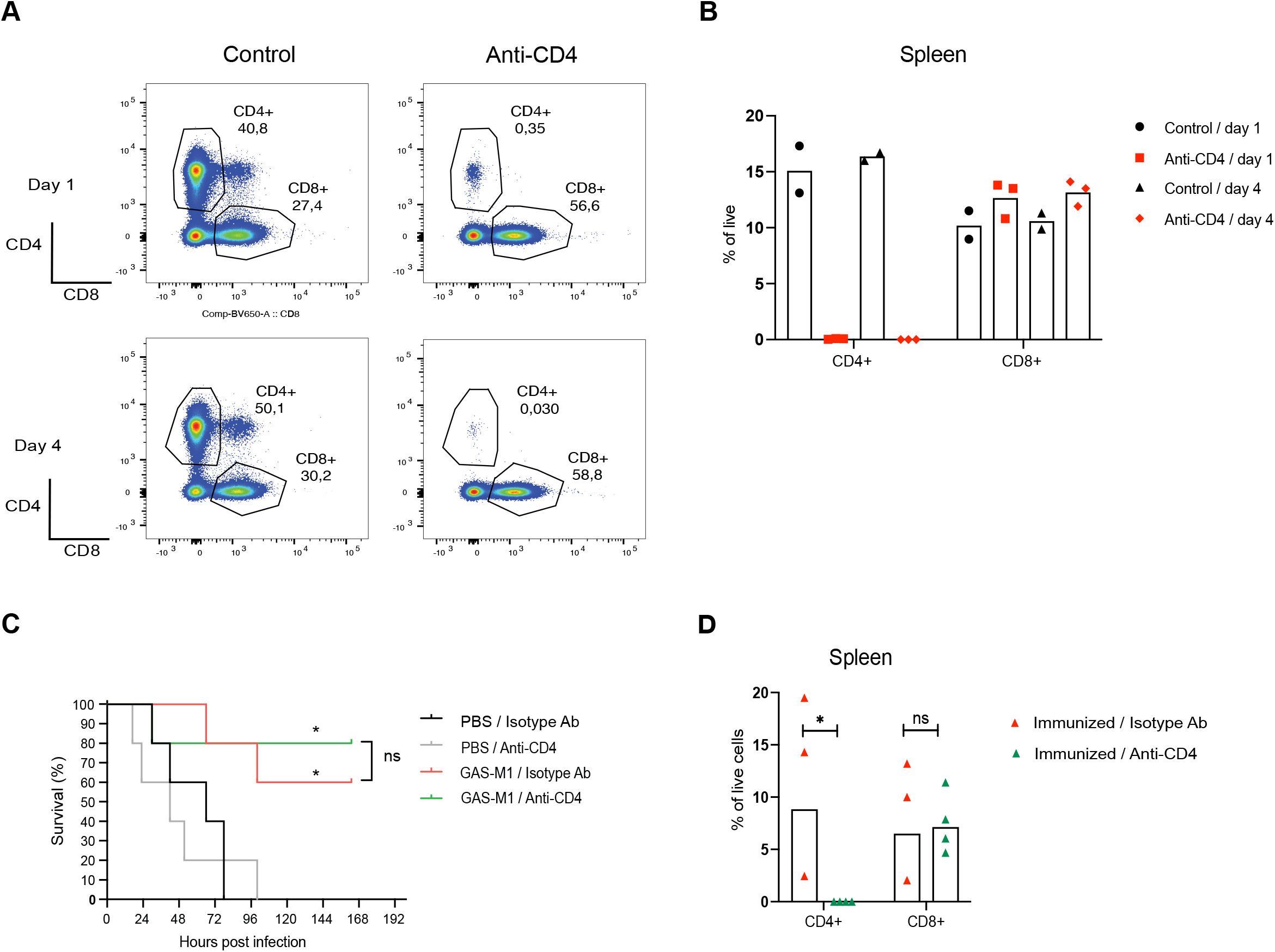
Depletion of CD4 T cells prior to lethal challenge does not affect survival of mice previously immunized with HK GAS-M1. (A-B) Validation of the CD4^+^ T cell depletion protocol. A control group of B6 mice were injected ip with a single dose of anti-CD4 or isotype control monoclonal antibodies and percentages of CD4^+^ and CD8^+^ T cells in spleen were determined by flow cytometry at day 1 and 4 after treatment. Results show gating strategy (A) and pooled results (B). (C-D) B6 mice were immunized ip with three doses of HK GAS-M1 with three weeks interval between each dose. Control B6 mice received 3 x PBS only. Depleting anti-CD4 monoclonal antibody was administered to half of the mice in each group 2 days before as well as 1 and 4 days after administration of a lethal dose GAS-M1 (n = 5 per group). Remaining mice in each group instead received an isotype control antibody by the same schedule as used for anti-CD4 treatment (n = 5 per group). Three weeks after the last immunization, all mice were infected ip with a lethal dose of GAS-M1 (10^8^ CFU) and then monitored for seven days. (C) Kaplan-Meier survival plot with log-rank tests with significant differences in relation to controls receiving PBS and isotype control antibody (PBS / isotype Ab) indicated. (D) Frequencies of CD4^+^ and CD8^+^ T cells in the spleen of immunized mice receiving anti-CD4 or control monoclonal antibodies and surviving the lethal dose until the experimental endpoint seven days post infection. *p<0.05, ns = not significant.

### Acquired protection in immunized mice is independent of adaptive immunity

The development of B and T cells requires the expression of RAG1 and RAG2, enzymes that mediate the rearrangement of the V(D)J segments of the B and T cell receptors [25]. In animals that lack *Rag1*, development is arrested at the pro-B/pro-T cell stage, leading to a severe deficiency in mature B and T cells [26]. To assess the overall contribution of adaptive immunity to protection in the ip HK GAS-M1 immunization model, *Rag1*-deficient and control B6 mice were immunized three times with HK GAS-M1 and then injected with a lethal dose of live GAS-M1 three weeks after last immunization. Strikingly, we observed no difference in survival between B6 and *Rag1*-KO, and both groups exhibited increased protection as compared to non-immunized mice (Fig 4A). As expected, we could not detect specific IgG or IgM in immunized *Rag1*-KO mice, whereas immunized B6 mice showed a considerable increase in both classes of antibodies (Fig 4B). Thus, our data suggest that adaptive immunity, *i.e.* B cells, T cells, and specific antibodies, are redundant for the protective immunity conferred by repeated ip immunization with HK GAS-M1.

**Fig 4.**
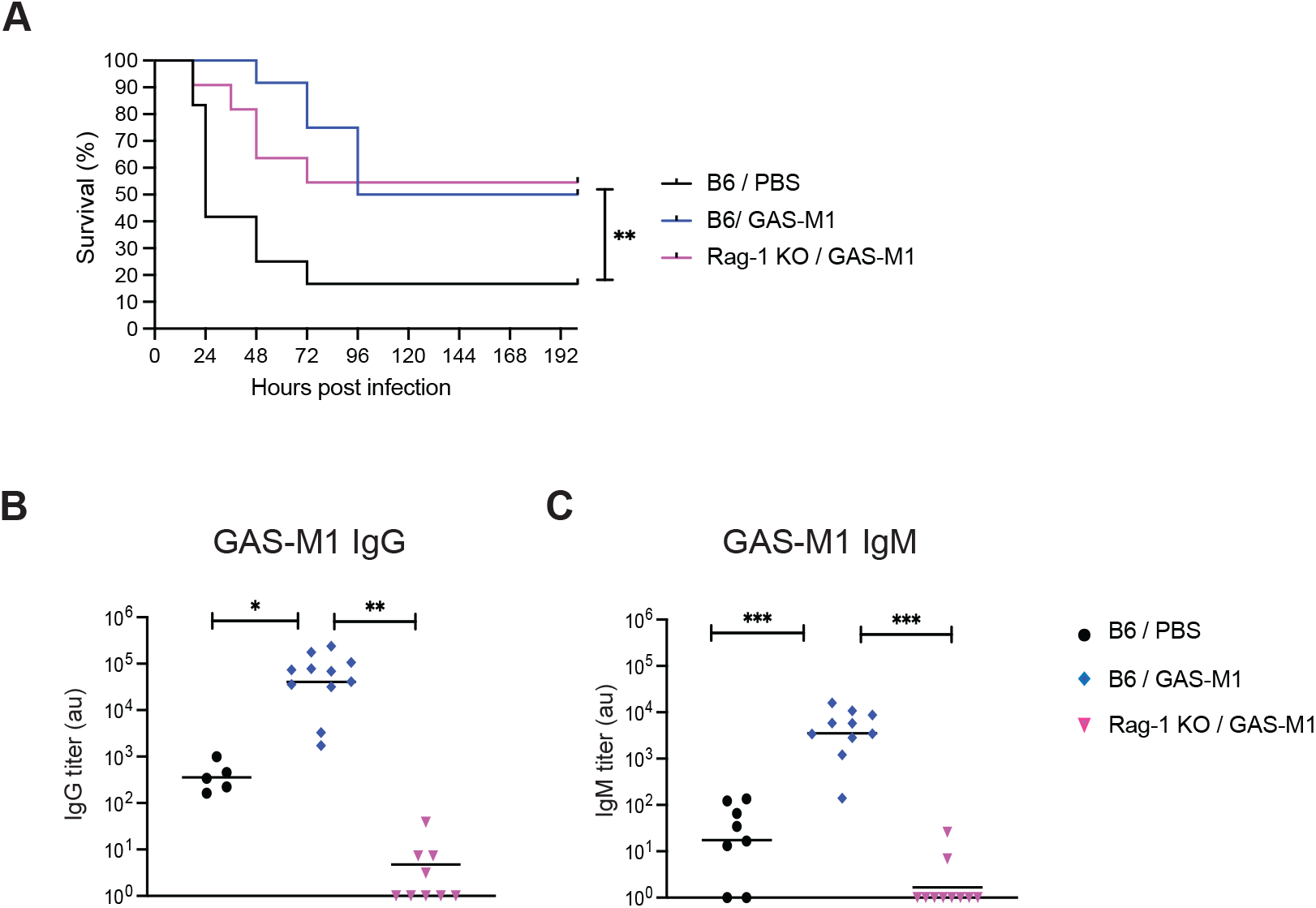
Acquired protection in immunized mice does not depend on B and T cells. B6 (n=10) and Rag1^-/-^ (n=10) mice were immunized ip three times with HK GAS-M1 with three weeks interval between each dose. Control B6 mice (n=10) received three doses of PBS only. Three weeks after the last immunization, all mice were infected ip with 10^8^ CFU GAS-M1 and monitored for eight days. Results are pooled from two separate experiments. (A) Kaplan-Meier survival plot with log-rank tests. (B) Levels of serum IgG and IgM against intact GAS-M1 three weeks after the last immunization and one day prior to administration of the lethal dose. Statistical significance was analyzed by one-way ANOVA with Tukey’s multiple comparisons test. *p<0.05, **p<0.01, ***p<0.001 and ****p<0.0001, ns = not significant.

### Selective increase in monocytes/macrophages and altered cytokine profile in immunized mice following lethal challenge with GAS-M1

As we found that protection in immunized mice was independent of adaptive immunity, we next analyzed the potential contribution of more acute innate responses. Immunized mice were again infected with a lethal dose of GAS-M1, but now euthanized 4 or 24 hours post infection and analyzed for bacterial dissemination as well as innate immune cell populations and cytokine profiles. Bacterial loads were similar in the peritoneal cavity, blood and spleen of immunized and non-immunized mice 4 hrs post-infection (Fig 5A). However, a significant increase in colony forming units (CFUs) were detected for all organs in non-immunized relative to immunized animals after 24 hrs (Fig 5A), demonstrating that early responses in immunized mice efficiently limit bacterial propagation. Focusing on the infiltration of immune cells into the peritoneal cavity of immunized and non-immunized mice, we observed no significant differences in numbers of neutrophils or eosinophils between the groups or time points (Fig 5B). Cells belonging to the monocyte/macrophage lineage, collectively identified as CD64^+^ cells lacking expression of lineage markers for T cells (CD3), B cells (B220) and NK cells (NK1.1), were relatively few 4 hrs post-infection but showed a dramatic increase in both groups 24 hours post-infection and, notably, was at this time point >10-fold higher in immunized vs non-immunized animals (Fig 5B and S2 Fig). In both steady-state and during inflammatory conditions, classical monocytes extravasate from the blood as Ly6C^+^ cells lacking expression of MHC class II (MHCII) and then undergo *in situ* differentiation into macrophages by a process where they first gain expression of MHCII (transitional monocytes; Ly6C^+^ MHCII^+^) and subsequently down-regulate Ly6C (macrophages; Ly6C^-^ MHCII^+^). A more detailed phenotypic analysis of the peritoneal monocytic cell population revealed that its pronounced increase in immunized animals 24 hrs post-infection was mostly driven by an accumulation of transitional Ly6C^+^ MHCII^+^ monocytes, with a minor contribution of fully differentiated Ly6C^-^ MHCII^+^ macrophages (Fig 5C). Collectively, these results indicate that a specific increase in monocytes and monocyte-derived macrophages occur after infection and that this is significantly enhanced by the ip immunization regimen.

Cytokine production was assessed in the peritoneal cavity, blood, and spleen of immunized and non-immunized mice (Fig. 5D and S3 Fig). For both groups of mice, the pro-inflammatory cytokines IL-1α, IL-1β and IL-6 were not detected 4 hrs post-infection but as expected increased in the peritoneal cavity of non-immunized mice at 24 hours. Curiously however, these cytokines were not elevated in immunized animals at this time point in spite of the increased presence of monocytes and macrophages (see Fig 5B and C) and protection against infection (see Fig 1C, 3C, and 4A). The chemokine CCL2 (MCP-1) is a key chemoattractant for monocytes [27]. In line with this function, CCL2 was selectively detected in immunized mice at 4 hrs (Fig 5D) – preceding the marked accumulation of monocytes/macrophages 24 hrs post infection in these animals (Fig 5B). In contrast, at 24 hrs post infection CCL2 was elevated in most of the non-immunized mice but could not be detected in the immunized group. A very similar pattern – *i.e.* an early production followed by decreasing levels in immunized mice and a late appearance in unimmunized animals – was observed for TNF-α, IFN-γ and IL-17A. The cytokine IL-5 was also selectively present in immunized mice at the 4 hrs time point but could not be detected at 24 hrs in any of the groups. Collectively, these results indicate that the immunization regimen promotes an accelerated production of CCL2, TNF-α IFN-γ, IL-17A and IL-5, while limiting the accumulation of the pro-inflammatory cytokines IL-1α, IL-1β and IL-6.

**Fig 5.**
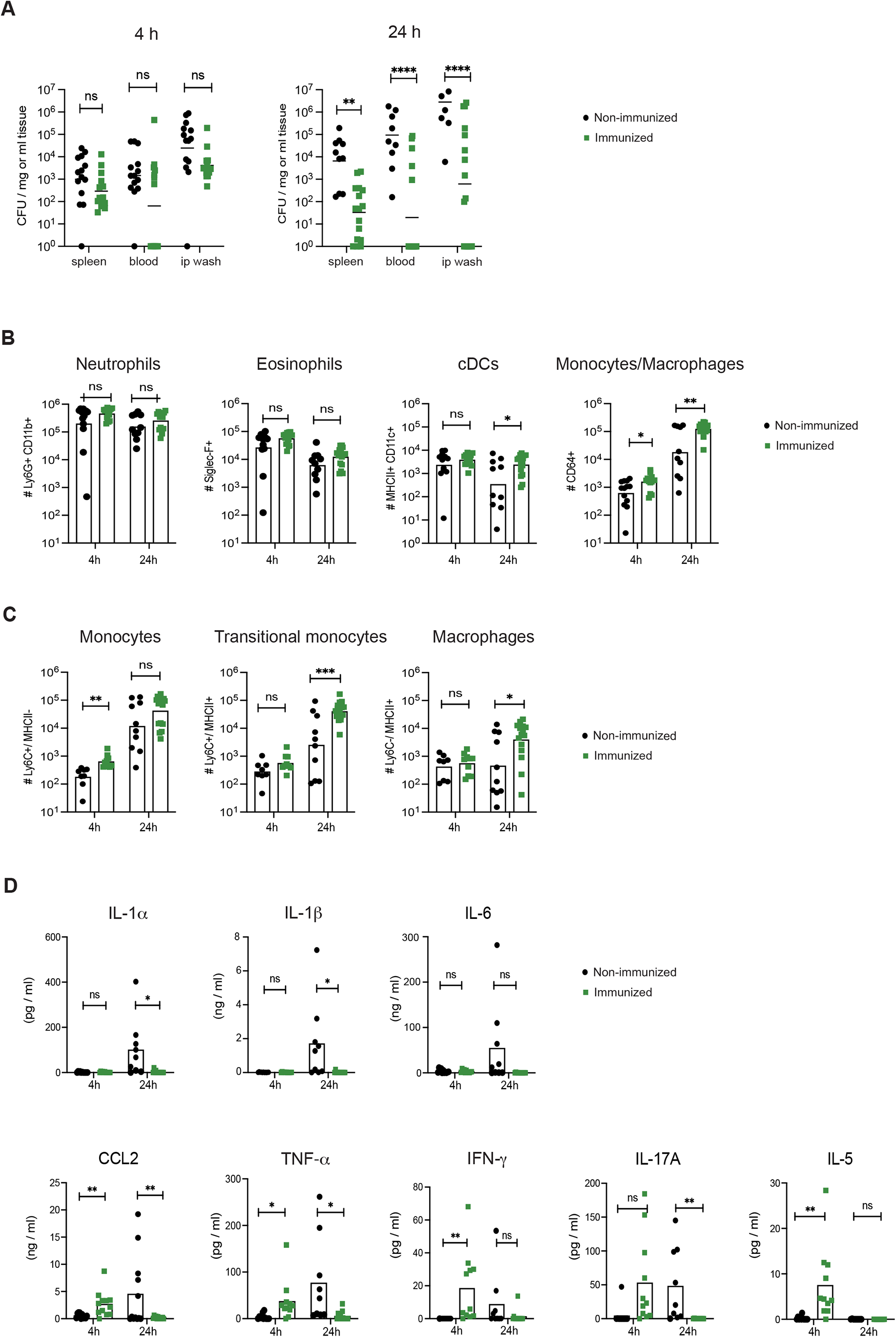
Reduced bacterial spread, elevated monocyte/macrophage numbers and an altered cytokine profile in peritoneum of immunized mice. B6 mice were immunized ip three times with HK GAS-M1 (n=31) or received 3 x PBS only (n=23). Three weeks after the final immunization, all mice were infected ip with 10^8^ CFU of GAS-M1. Mice were euthanized either 4 or 24 hours post-infection for analysis. (A) Bacterial loads in the spleen, blood and peritoneal cavity (ip wash). (B) Numbers of neutrophils, eosinophils, dendritic cells and CD64^+^ cells of the monocyte/macrophage lineage in the peritoneal cavity (ip wash) were determined by flow cytometry and total cell count. (C) Subset analysis of the peritoneal CD64^+^ cells by flow cytometry, showing numbers of Ly6C^+^MHCII^-^ monocytes, Ly6C^+^MHCII^+^ transitional monocytes and Ly6C^-^MHCII^+^ macrophages. (D) Concentration of CCL2 and indicated cytokines in the peritoneal cavity (ip wash). Pooled results from three separate experiments are shown. Statistical significance was analyzed by unpaired t-test comparing the mean of each group. *p<0.05, **p<0.01, ***p<0.001 and ****p<0.0001, ns = not significant.

### The immunization-induced protection against GAS infection is dependent on macrophages

As we observed a significant and selective increase in monocytes/macrophages in the peritoneal cavity after ip infection of immunized mice, we hypothesized that these cell populations may play an essential part in the induced protection in immunized animals. Macrophages can be specifically depleted using liposome-encapsulated clodronate [28]. Injected liposomes are taken up through endo-or phagocytosis, and after vacuolar fusion with lysosomes the clodronate is released and induces apoptosis of the cell [29]. In clodronate treated mice, macrophages were effectively and specifically depleted, as determined 3 days after liposome injection (Fig 6A and S4 Fig). We then treated immunized mice with ip injections of clodronate 3 days prior to and one day post lethal dose to ensure absence of macrophages throughout the infection, control mice received liposomes containing PBS only (Fig 6B). While control mice remained protected, clodronate treated animals exhibited significantly reduced ability to withstand infection, suggesting that macrophages comprise an essential part of the protective immune mechanism established by the ip immunization regimen.

**Fig 6.**
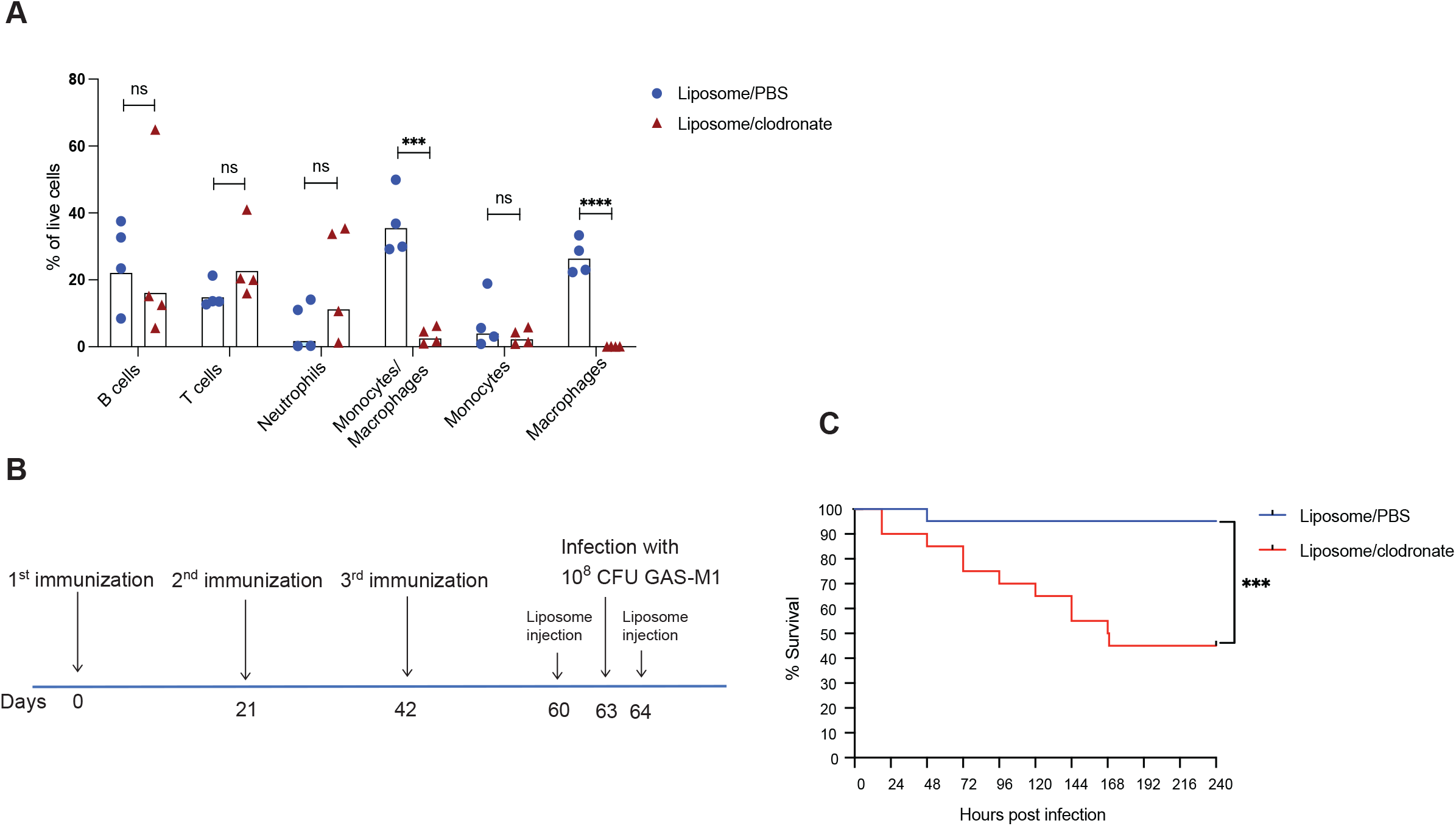
Macrophages are required for protection against GAS-M1 infection in immunized mice. (A) Assessment of macrophage-depletion by clodronate-liposomes. Mice were injected ip with clodronate-liposomes (2.5 mg/mouse) or PBS-liposomes (2.5 mg/mouse) and cellular composition in the peritoneal cavity was assessed by flow cytometry 3 days later. Results show percentage of indicated subset among total viable cells and are pooled from two separate tests. Statistical significance was analyzed by unpaired t-test comparing the mean of each group. (B-C) B6 mice (n=21) were immunized three times with HK GAS-M1and then challenged with lethal infection three weeks after last immunization. One group received an ip injection of clodronate-liposomes both three days before and one day after lethal infection while the control group received PBS-liposome injections at the same time points. Mice were monitored for 10 days post-infection. (B) Experimental layout. (**C**) Kaplan-Meier survival plot with log-rank tests. *p<0.05, **p<0.01, ***p<0.001 and ****p<0.0001.

### IFN-γ is a key determinant of protection against GAS infection in immunized mice

In immune mechanisms of protection against infection, activated Th1 cells are a major source of the cytokine IFN-γ, potentiating macrophage capacity to eliminate intracellular pathogens [30]. IFN-γ has however also been implicated in the more acute innate defenses against infection with extracellular pathogens, including GAS [31, 32]. Given the rapid increase in peritoneal IFN-γ levels in immunized mice following administration of the lethal dose (see Fig 5D), we assessed protection in IFN-γ deficient mice and found that in the complete absence of IFN-γ, even immunized animals succumbed to GAS-M1 infection (Fig 7), suggesting a significant role for this cytokine in immunization-induced protective immunity.

**Fig 7.**
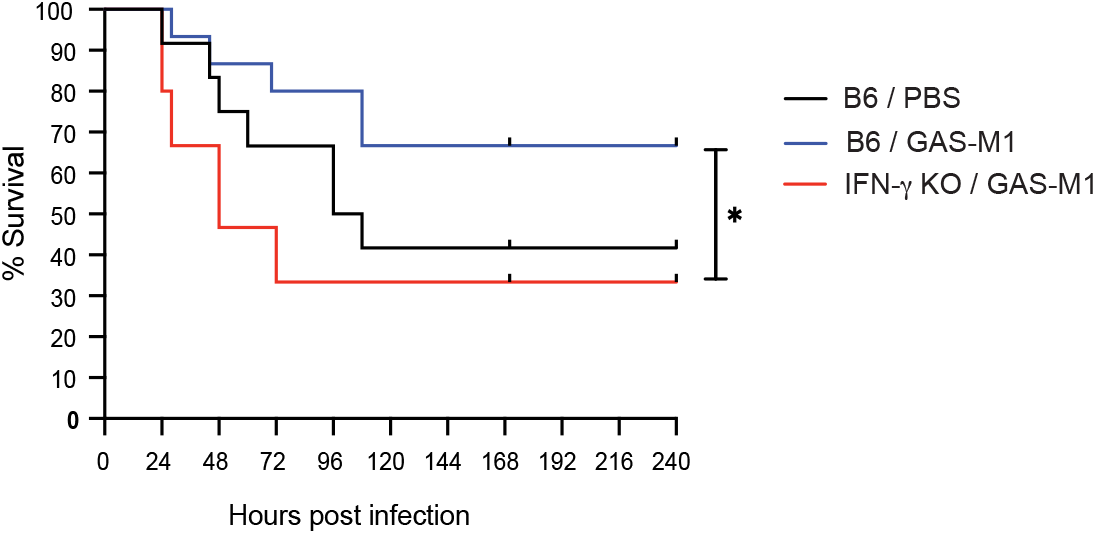
IFN-γ is required for protection against GAS-M1 infection in immunized mice. B6 (n=15) and IFN-γ knock-out (n=15) mice were immunized ip three times with HK GAS-M1 with three weeks between each dose. Control B6 mice received three doses of PBS only (n=12). Three weeks after the last immunization, all mice were infected ip with 10^8^ CFU GAS-M1 and monitored for ten days. Results are pooled from two separate experiments. Kaplan-Meier survival plot with log-rank tests is shown. *p<0.05

## Discussion

In this study, we demonstrate that recurring injections with HK wt GAS-M1 into the murine peritoneal cavity generates protective immunity. Further we show that although the ip immunization regimen did induce production of GAS-specific IgG, several lines of evidence excluded the involvement of adaptive immune mechanisms such as GC development, CD4 T cells, and antibodies, in the protective response conferred by the immunization. Protection was instead dependent on the presence of macrophages during the infection phase, and surprisingly on the cytokine IFN-γ although CD4 T cells were not required (Th1 cells are commonly a major source of IFN-γ in recalled immune responses). These findings suggest that protection in our model may rather be driven by trained innate immune cell populations.

The trained immunity phenomenon was recognized with the realization that live attenuated vaccines, most notably BCG, in addition to the intended protection against the vaccine pathogen, conferred heterologous protection leading to reduced incidence of several common diseases in BCG-vaccinated children [33, 34]. This protection has since been found to be driven by metabolic and epigenetic changes in innate immune cell precursors in the bone marrow (central trained immunity) and in differentiated circulating or tissue resident cells (peripheral trained immunity), leading to altered response quality and quantity upon secondary challenges [8]. Much effort has been put into deciphering the molecular signatures defining different forms of trained immunity, but little is still known about the specific requirements for when trained immunity is triggered *in vivo*. Our previous work in a model of subcutaneous (sc) immunization showed that protective immunity was not generated against wt GAS, but required the insertion of a dominant CD4 T-cell epitope in a bacterial surface protein, and was dependent on canonical adaptive immunity [35]. The disparity in how protection is generated in our sc and ip models clearly highlights the radical difference in immune responses and subsequent development of both adaptive and innate protective immunity in distinct tissues. Interestingly, related observations have been made in experimental models of BCG immunizations, where skin injections appear to only induce trained immunity if the inoculum dose is sufficiently low [36]. These observations might be providing intriguing clues into the requirements for what triggers trained immunity *in vivo*.

Little is known about the ability of many common bacterial species other than Mycobacteria to induce trained immunity. Of particular interest for our work on GAS, a handful of studies using other Gram-positive encapsulated bacteria suggest *e.g.* that *Staphylococcus aureus* sc injections induce trained immunity that protect against subsequent local skin infection [37–39]. Further, a recent study suggests that intranasal infection in mice using *Streptococcus pneumoniae* elicits specific NK-cell mediated protection against a subsequent lethal bacterial dose [40]. In contrast, in a study of trained immunity also employing intranasal immunizations, protection against subsequent infection was induced by several Gram-negative bacterial species, but not by *S. aureus* or *S. pneumoniae* [41]. Of note, the immunomodulant compound OM-85 is a bacterial lysate mix containing lyophilized fragments of 21 common bacterial pathogens, including GAS, *S. aureus* and *S. pneumoniae*, and is used clinically to treat children suffering from recurrent lower respiratory tract infections [42, 43]. The exact mechanism(s) of action of this treatment regimen has not yet been elucidated, but as protection is evident also for viral pathogens, the mechanism of protection is believed to at least partially be attributed to induced trained immunity.

Studies of trained innate immune cells *in vitro* have shown that different populations present with distinct reaction phenotypes, *e.g.* trained monocytes and macrophages produce elevated levels of IL-6, while trained NK cells often present with increased production of IFN-γ [8]. In a whole organism context *in vivo* these cell-specific distinctions are not as easily discerned, and in our model the source of different cytokines can at this stage only be presumed. Indeed, we were curious to observe that protection was dependent on macrophages in immunized mice, although these animals did not present with an obvious total cytokine profile agreeing with what is expected from re-called trained macrophages. Instead, we detected accelerated and increased levels of IFN-γ, IL-5 and IL-17, combined with an apparent absence of classical pro-inflammatory cytokines, including IL-6, upon infection of immunized mice. While the cellular source(s) of these cytokines remains elusive, the cytokine profile imprinted by the ip immunizations with HK wt GAS seems to be in line with a possible involvement of ILC populations, including IFN-γ producing memory-like NK cells and possibly trained ILC2 and ILC3 producing IL-5 and IL-17, rather than myeloid subsets [44, 45]. IFN-γ producing tissue resident NK cells are indeed present in the peritoneal cavity [46] and it seems possible that NK cell-derived IFN-γ increases the bactericidal capacity of macrophages in the immunized mice, thus restricting bacterial growth and dissemination, and promoting survival after infection [47, 48]. Our results also show elevated and accelerated CCL2 production in the immunized mice, and given the central role for CCL2 in regulating monocyte migration and function [27], it seems likely that this chemokine is at least partially responsible for the enhanced recruitment of monocytic cells to the peritoneal cavity of immunized mice following the challenge with live bacteria. How the current immunization regimen imprints altered magnitude and kinetics of CCL2 production remains to be determined. Finally, we are hesitant to firmly conclude that we are observing a *de facto* inhibited production of pro-inflammatory cytokines in infected immunized mice. As it is likely that the reduced bacterial proliferation in these animals lead to weaker stimulation for proinflammatory cytokine production, and possible that augmented consumption comes with increased levels of incoming CD64^+^ myeloid cells, this question should be addressed in a context that allows for sufficient stratification of data.

In conclusion, our results reveal a novel aspect of protective immunity generated against GAS that depends on IFN-γ, involves enhanced recruitment of monocytes and their local differentiation into macrophages, and occurs independently of adaptive memory responses. Future mechanistic studies addressing training of innate immune cell populations as well as the origin and function of distinct mediators in immunized mice should shed further light on the role and requirements of protective innate responses in GAS infections and vaccination strategies.

## Materials and methods

### Bacterial strains and culture conditions

The *S. pyogenes* strain 90-226 is an M1 serotype strain originally isolated from a sepsis patient [49] and is referred to as GAS-M1 in this study. Bacterial cultures were grown over night without shaking in Todd-Hewitt broth supplemented with 0.2% yeast extract (THY) in 5% CO2 at 37^°^C, and re-inoculated and grown until they reached optical density (OD) of 0.8-1.

### Mice

Female C57Bl/6 (wildtype, [wt]) mice were purchased from Taconic and IFN-γ-KO mice (B6.129S7-Ifng^tm1Ts^/J) were originally from The Jackson Laboratory. The CD4-Cre x Bcl6^fl/fl^ strain was generated by crossing CD4-Cre (B6.CgTg(Cd4-cre)1Cwi/BfluJ) and Bcl6^fl/fl^ (B6.129S(FVB)Bcl6 ^tml.1Dent^/J) mice. Animals with the CD4-Cre x Bcl6^fl/fl^ genotype do not develop T follicular helper (Tfh) cells, as they lack the Tfh master transcription factor Bcl6 in CD4^+^ T-cells. Rag1-KO mice on a B6 background were originally from The Jackson Laboratory. For experiments using wt mice only, 8-week-old females were used while genetically modified mice were both female and male, aged 8 – 12 weeks. All animals were bred (except B6) and maintained at the animal facility at the Lund University Biomedical Center, and experiments were performed in accordance with protocols approved by the Lund/Malmö Animal Ethics Committee.

### Immunizations and infections

To generate heat killed (HK) or formalin fixed (FF) bacteria, log phase cultures were washed with PBS and incubated at 60°C for two hours, or fixed with 1% formalin to achieve complete killing. Killing was confirmed by blood agar plating. Suspensions of killed bacteria were diluted to the appropriate concentration and stored at -20°C (HK) or 4 °C (FF). Mice were injected with 10^8^ HK or FF GAS-M1 once, or three times ip at three-week intervals and bled 19 days after each injection. For protection experiments, a lethal dose of 10^8^ CFU live GAS-M1 was administered ip three weeks after the last immunization injection, after which the cages were blinded to prevent any bias in determining moribundity.

### Serum antibody titer measurements

To assess antibody titers in mouse serum, HiBond ELISA plates were coated with 10^7^ CFU of GAS-M1 bacteria in a buffer containing PBS and 0.05% Sodium azide. These coated plates were incubated overnight at 4°C and on the following day incubated with serially diluted serum samples obtained from both immunized and non-immunized mice. Captured antibodies were detected using biotinylated goat anti-mouse IgG (Southern Biotech, CAT: 1036-08) and streptavidin-HPR (BioLegend, Cat 405210). All samples were run in duplicate and presented values represent the average of duplicates. BSA-coated wells served as negative controls.

### Detection of bacterial counts, immune cells, and cytokines in different organs of infected mice

On the selected time points post infection, animals were sacrificed, and 5 ml sterile PBS was injected into the peritoneal cavity. The abdominal area was massaged briefly and the ip wash was extracted using a syringe and kept on ice. The total number of recovered white blood cells was determined in a Sysmex hematology analyzer (Sysmex), and bacterial numbers were determined by blood agar plate culture and manual counting. Blood was collected from the heart and mixed with 0.5 M EDTA to prevent coagulation. The spleen was removed, weighed, and homogenized using a 70 μm cell strainer. Serial dilutions of blood and spleen samples were cultured on blood agar plates for determination of bacterial loads. Immune cells were identified in ip wash, blood or homogenized spleen by flow cytometry, using the following reagents and fluorochrome conjugated antibodies: Live/dead fixable APC-Cy7 (Invitrogen, L34975), Anti-CD3-AF700 (BD, 17A2), Anti-NK-1.1-AF700 (BioLegend, PK136), Anti-CD19-AF700 (BioLegend, 6D5), Anti-CD11b-BV605 (BioLegend, M1/70), Anti-CD11c-PE-CY7 (Invitrogen, N418), Anti-MHC-II-BV421 (BioLegend, M5/114.15.2), Anti-Gr-1-BV510 (BioLegend, RB6-8C5), Anti-Ly6C-PerCP-Cy5.5 (BioLegend, HK1.4), Anti-CD64-AF647 (BD Pharmigen, X54-5/7.1), Anti-Siglec-F-PE (BD Pharmigen, E50-2440). Cytokine concentrations were measured using CBA (single-color antibodies; BD Biosciences).

### Depletion of T cells and macrophages

For T cell-depletion experiments, mice were injected ip with a depleting anti-CD4 antibody (BioxCell, BE0003-1) two days before infection, and on post infection days +1 and +4. Control mice received isotype control antibody (BioxCell, BE0090) only. Spleen cells were recovered as described above and T cells were analyzed by BD **^®^** LSR II Flow Cytometer using the following reagents and fluorochrome conjugated antibodies: Live/dead fixable APC-Cy7 (Invitrogen, L34975), Anti-CD3-BV605 (BD, 17A2), Anti-CD4-FITC (eBioscience, RM4-5), Anti-CD8-BV650 (BD, 53-6.7).

To deplete macrophages, immunized mice were injected ip with 500 μl clodronate liposomes (5 mg/ml; LIPOSOMA BV, NL) three days prior to and one day post infection. Immunized control mice received PBS liposomes at the same time points. To identify cells in the peritoneum (ip wash, as described above), the following reagents/antibodies were used: Anti-CD3-AF700 (BioLegend, 17A2), Anti-CD11b-BV786 (BD Bioscience, M1/70), Anti-CD19-BV711 (BioLegend, 6D5), Anti-CD64-APC (BD Bioscience, X54-5/7.1), Anti-F4/80-BV421 (Biolegend, BM8), Anti-Ly6C-FITC (BioLegend, HK1.4), Anti-Ly6G-PE (BioLegend,1A8), and live/dead PI-PE-CF594 (Invitrogen, CAT P3566).

### Statistical analysis

Data were analyzed using Prism version 8.0 (GraphPad Software). Analysis of statistical significance was performed using one-way ANOVA with Tukey’s multiple comparisons test for three or more groups, or t-test for two unpaired groups. Comparison of survival curves was done by Kaplan-Meier survival test with Log-rank (Mantel-Cox). Differences were considered significant when p ≤ 0.05 (*p≤0.05, **p<0.01 and ***p<0.001).

## Acknowledgements

We are grateful to Dr Patrick Cleary and Dr Thamotharampillai Dileepan for kindly providing the 90-226 GAS-M1 strain used in this work.

## Supporting information

**S1 Fig.**
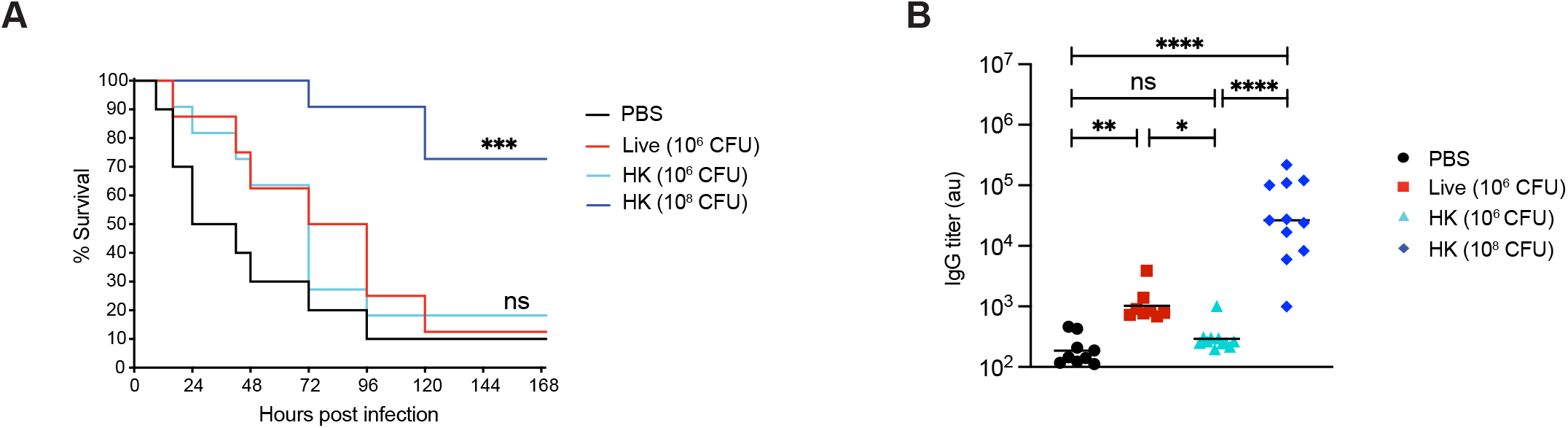
Protection conferred by repeated intraperitoneal immunization with HK GAS-M1 requires a high dose of HK bacteria. B6 mice (n = 10-11 per group) were immunized ip with three doses of GAS-M1 with three weeks intervals of live (10^6^ CFU; highest non-pathological dose possible), FF (10^8^ CFU) or HK (10^6^ and 10^8^ CFU, respectively) bacteria. Control B6 mice received PBS only. Three weeks after the third immunization, mice were challenged ip with a lethal dose of GAS-M1 (10^8^ CFU) and monitored for 7 days. (A) Kaplan-Meier survival plot with log-rank tests. (B) Serum level of IgG against intact GAS-M1 19 days after the third immunization and prior to administration of the lethal dose. Statistical significance was analyzed by one-way ANOVA with Tukey’s multiple comparisons test. **p<0.01, ***p<0.001 and ****p<0.0001. ns = not significant.

**S2 Fig.**
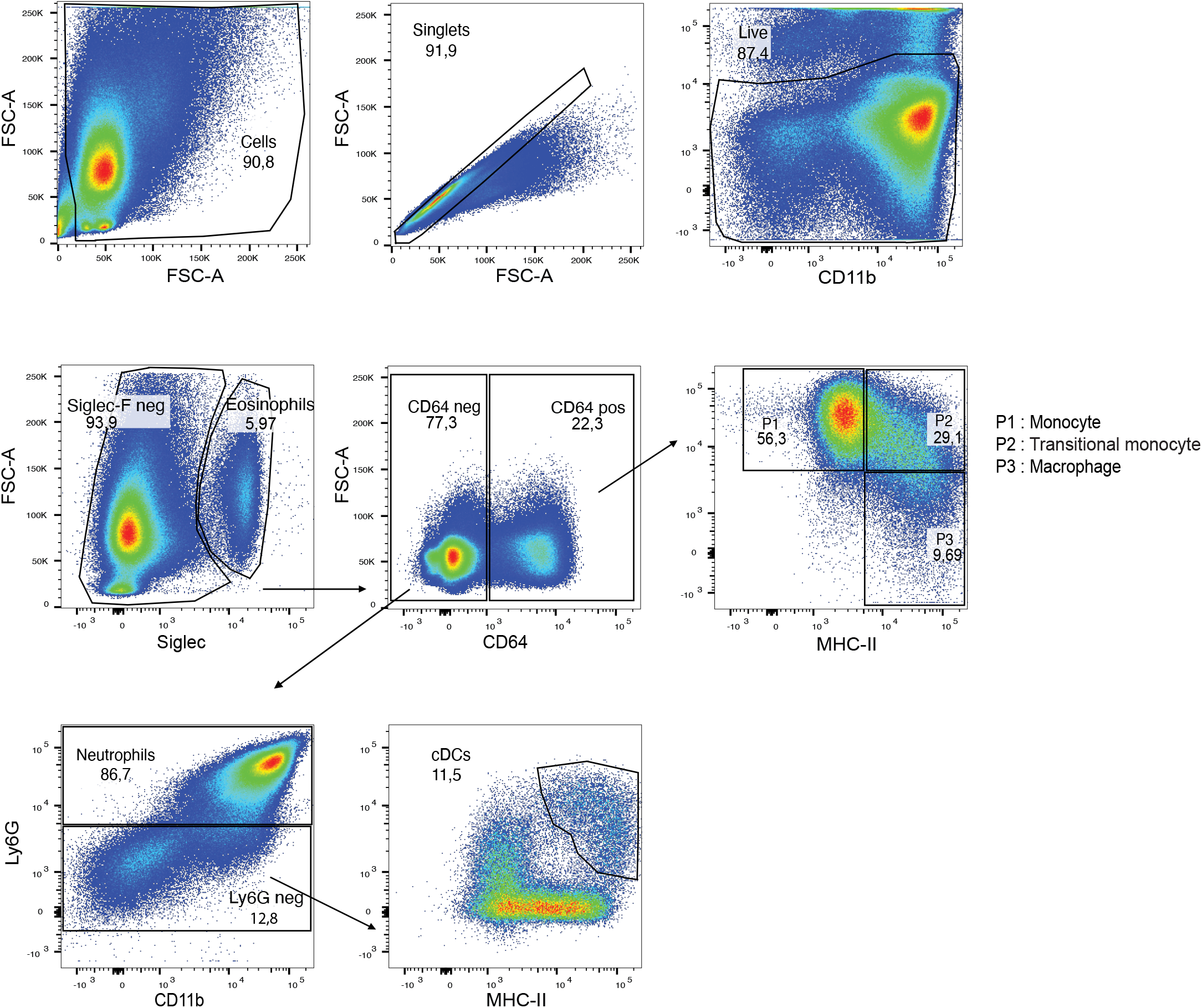
Flow cytometry analysis of peritoneal leukocyte subsets in immunized and non-immunized mice after lethal GAS-M1 challenge. Gating strategy used for phenotypic definition of eosinophils, neutrophils, dendritic cells, total cells of the monocyte/macrophage lineage, monocytes, transitional monocytes, and macrophages in the peritoneal cavity of mice described in Figure 5B and C.

**S3 Fig.**
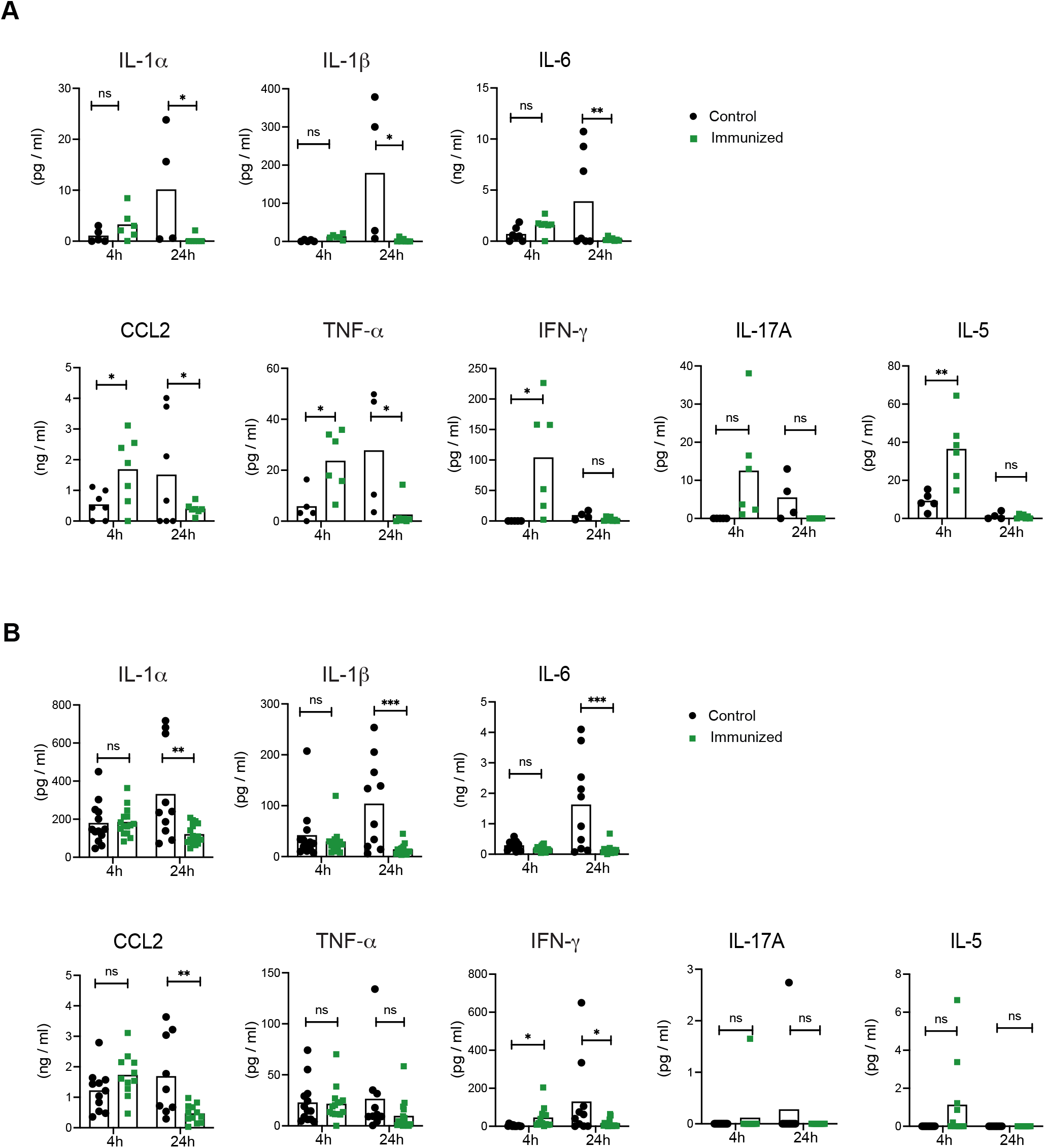
Distinct cytokine profiles in blood and spleen from immunized and non-immunized mice. Immunized and non-immunized mice were infected with 10^8^ CFU GAS-M1 and sacrificed after 4 or 24 hours as described in Figure 5D. Cytokine levels in serum (A) and homogenized spleen (B) of individual mice. *p<0.05, **p<0.01 and ***p<0.001. ns = not significant. Related to Figure 5D.

**S4 Fig.**
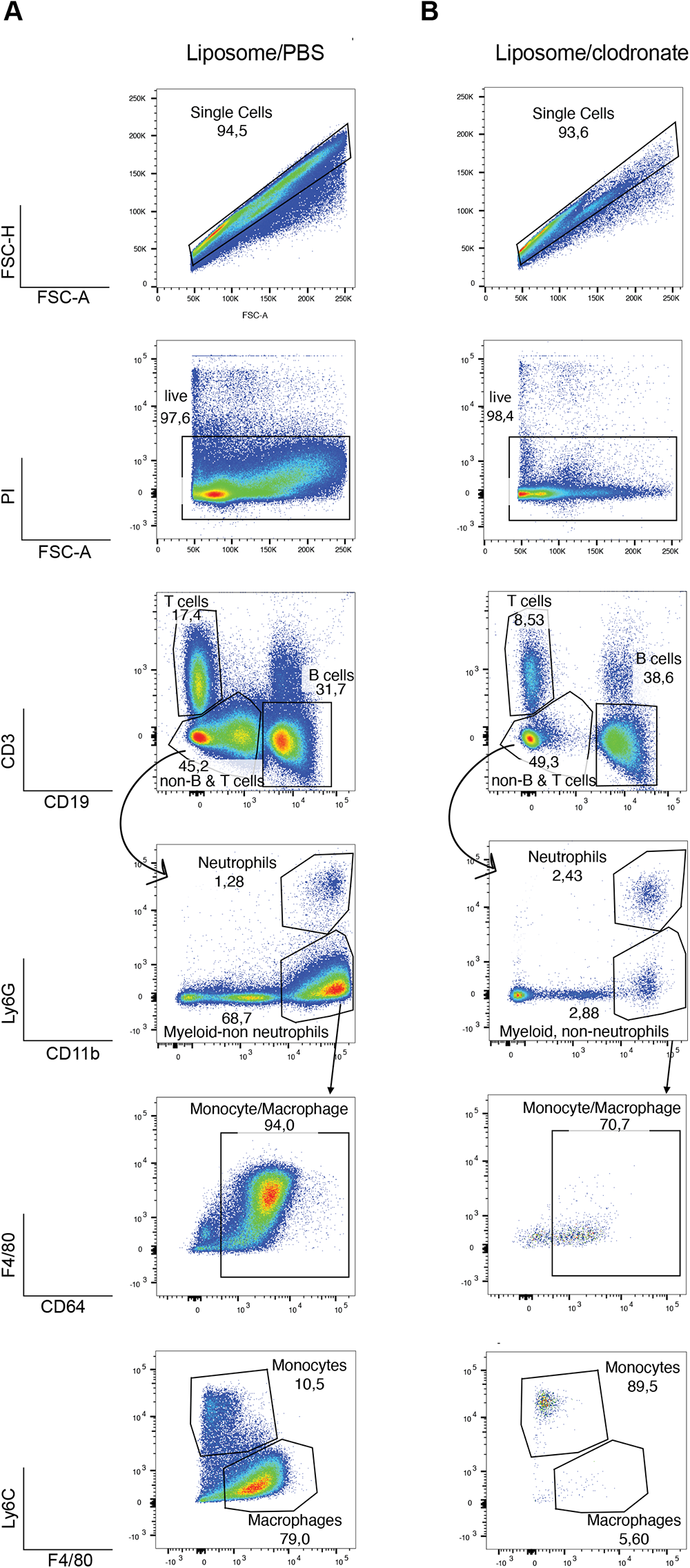
Flow cytometry analysis of peritoneal leukocyte subsets following treatment with clodronate liposomes. Gating strategy used for phenotypic definition of T cells, B cells, neutrophils, total cells of the monocyte/macrophage lineage, monocytes and macrophages in the peritoneal cavity of mice treated with liposomes containing PBS (A) or clodronate (B). Related to Figure 6.

